# LSTrAP-Cloud: A User-friendly Cloud Computing Pipeline to Infer Co-functional and Regulatory Networks

**DOI:** 10.1101/2020.03.11.986794

**Authors:** Qiao Wen Tan, William Goh, Marek Mutwil

## Abstract

As genomes become more and more available, gene function prediction presents itself as one of the major hurdles in our quest to extract meaningful information on the biological processes genes participate in. In order to facilitate gene function prediction, we show how our user-friendly pipeline, Large-Scale Transcriptomic Analysis Pipeline in Cloud (LSTrAP-Cloud), can be useful in helping biologists make a shortlist of genes that they might be interested in. LSTrAP-Cloud is based on Google Colaboratory and provides user-friendly tools that process and quality-control RNA sequencing data streamed from the European Sequencing Archive. LSTRAP-Cloud outputs a gene co-expression network that can be used to identify functionally related genes for any organism with a sequenced genome and publicly available RNA sequencing data. Here, we used the biosynthesis pathway of *Nicotiana tabacum* as a case study to demonstrate how enzymes, transporters and transcription factors involved in the synthesis, transport and regulation of nicotine can be identified using our pipeline.

## 1. Introduction

Genome sequencing and assembly are becoming more accessible in terms of cost and computational resources required due to the advances in technology and algorithms [1]. However, elucidating gene function is necessary to extract meaningful knowledge from genomes. Despite extensive efforts over the decades, only 12% of genes have been characterised in the most studied model plant Arabidopsis thaliana [2]. This is because gene characterisation is a time and labour intensive process hindered by various obstacles such as the lethality of mutants involving essential genes, or conversely, no observable mutant phenotype due to functional redundancy caused by large gene families [3–7]. Evidently, unguided experimental characterization of all genes is not feasible and consequently, computational gene function prediction studies are conducted to meet this challenge (reviewed in Rhee and Mutwil). To this end, newly sequenced genomes are mostly annotated using sequence similarity approaches, which annotate novel genes based on the sequence similarity to characterized genes. While sequence similarity analysis gives a quick overview of gene functions in a new genome, it has its limitations as genes can i) have multiple functions, ii) sub- or neo-functionalise via gene duplications or iii) have no sequence similarity to characterized genes. Most importantly, sequence similarity approaches often cannot reveal which genes work together in a novel biological process, for example, a specialized metabolic pathway. Clearly, classical approaches to gene function prediction are powerful but require other methods to complement it [3].

In order to associate genes to pathways, one has to consider the expression of genes at the level of organs, tissues and even cells. With the increasing availability of RNA sequencing (RNA-seq) data, it is now possible to study genes from the perspective of their expression [3,8]. Genes that have similar expression profiles across different organs, developmental stages, time of the day, biotic and abiotic stress conditions tend to be functionally related [3,6,8–11]. These transcriptionally coordinated (co-expressed) genes can be revealed by analyzing transcriptomic data stemming from microarrays or RNA-seq data. In turn, co-expressed genes can be represented as nodes connected by edges (links) in co-expression networks, which can be mined for groups of highly connected genes (modules, clusters) that are likely to be involved in the same biological process. Co-expression networks have become a popular tool to elucidate the function of genes, their related biological processes and gene regulatory landscapes. Genes involved in a wide range of processes including cellular processes [12–14], transcriptional regulation [15], physiological response to the environment and stress [9,16], plant viability and biosynthesis of metabolites [17–21] have been elucidated using co-expression analyses.

The amount of gene expression data has grown tremendously over the decade, with more than 1000-fold increase in nucleotide bases on NCBI Sequence Read Archive (SRA), from 11TB to 12 PB at the start of 2010 and 2020, respectively. Analysing this data would have been unthinkable a decade ago, due to limitations in software used to estimate gene expression from RNA-seq data. However, drastic improvements in software used to estimate gene expression from RNA-seq data, such as kallisto [22] and salmon [23], have made this task possible within reasonable time on a typical desktop or even a Raspberry Pi-like miniature computer [24]. Furthermore, multiple user-friendly pipelines are an invaluable resource both for experts and non-bioinformaticians, to which pipelines such as UTAP [25], CURSE [26], LSTrAP-Lite [24] and LSTrAP [27] are made publicly available. However, all these resources typically require complex installation or a linux environment.

The introduction of cloud computing has provided alternatives to how data can be managed, stored and processed. Google colaboratory (colab), a Jupyter notebook environment was launched in 2017 and it allows users to write and execute python code on Google’s cloud servers through their browser (https://colab.research.google.com/). The use of the colab platform for RNA-seq analysis was demonstrated by Melsted et. al., 2019, who implemented the workflow of pre-processing single cell transcriptomic data. The choice of colab greatly improves user friendliness with the clean layout of jupyter notebooks, a graphical interface that is more friendly for biologists compared to the typical linux terminal, and problems associated with installation in different local environments. Collaboration between scientists is also easily established as notebooks can be saved to Google Drive and Github and deployed on a new computer within a minute. Most importantly, colab provides the computing power to perform RNA-seq analysis for free, allowing the users to run computationally heavy calculations on any computer, tablet or cell phone.

These advances and resources have prompted us to showcase the use of colab as a seamless and user friendly interface for large scale transcriptomic analysis. We illustrate this by using RNA-seq data of the model plant *Nicotiana tabacum* to dissect the biosynthesis of nicotine by co-expression network analysis. The presented pipeline, Large-Scale Transcriptome Analysis Pipeline in Cloud (LSTrAP-Cloud), is available from https://github.com/tqiaowen/LSTrAP-Cloud and can be easily applied to other pathways and organisms.

## 2. Materials and Methods

### 2.1. Streaming RNA sequencing data

The pipeline (Figure 1) was implemented on Google Colaboratory and consists of two jupyter notebooks that are available on https://github.com/tqiaowen/LSTrAP-Cloud. RNA sequencing experiments (Table S1) were streamed as fastq files from the European Nucleotide Archive (ENA) [28] using curl v7.58.0 with the parameters “-L -r 0-1000000000 -m 600 –speed-limit 1000000 –speed-time 30”, which allows curl to be redirected to updated address, retrieve first 1 billion bytes (953 megabytes) of the fastq file, run for maximum duration of 600 seconds and abort the download if speed drops below 1 million bytes per second (1 Mb/s), respectively. Files ending with “_1.fastq.gz” and “.fastq.gz” were downloaded for paired and single library layouts respectively. The streamed data from curl was piped to kallisto quant v0.46.0 [22] with the parameters “–single -l 200 -s 20 -t 2” (single end experiment with read length of 200 bp, standard deviation of 20 and to run with two threads) and mapped against the kallisto index of coding sequences (CDS) of *Nicotiana tabacum* [29]. CDS Nitab-v4.5_cDNA_Edwards2017.fasta was obtained from SolGenomics [30] and used to generate kallisto index with default parameters. A total of 1049 out of 1060 experiments were processed successfully. Seven files were not found and four files had unacceptable download speed among the files that were not processed successfully (Table S2).

**Figure 1.**
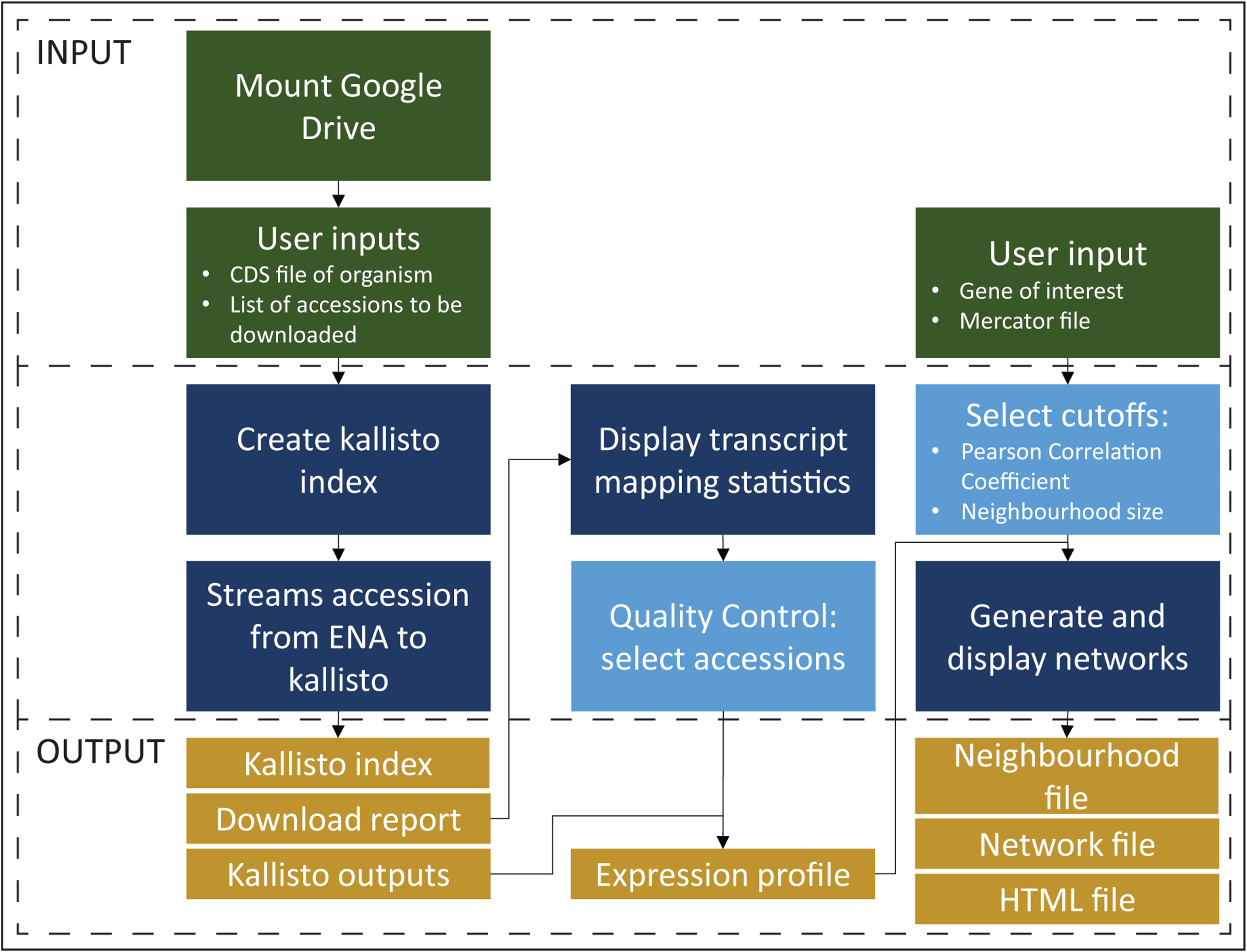
Schematic of LSTrAP-Cloud pipeline used for streaming and mapping of RNA sequencing data, and for the generation of co-expression network based on the gene of interest.

### 2.2. Constructing co-expression networks

Co-expression network of the gene of interest, *Nitab4.5_0000884g0010.1*, was obtained by including a maximum of 50 other genes with a Pearson Correlation Coefficient of at least 0.7 against the gene of interest. The co-expression network of *Nitab4.5_0000884g0010.1* was visualised on colab using Cytoscape.js v3.9.4 [31]. The shapes and colours of the genes were assigned according to the major Mapman bin classification obtained from Mercator4 v2.0 [32] with the *N. tabacum* Nitab-v4.5_cDNA_Edwards2017.fasta CDS (Table S3). The co-expression neighbourhood of the gene of interest is also displayed at the end of the colab notebook.

To generate Figure 3, The co-expression network of *Nitab4.5_0000884g0010.1* was downloaded from colab as a JSON file and modified in Cytoscape desktop v3.7.1 (Table S4). For brevity, only transporters, transcription factors and genes involved in nicotine biosynthesis are shown, but the network containing all 50 genes is available (Figure S1).

**Figure 2.**
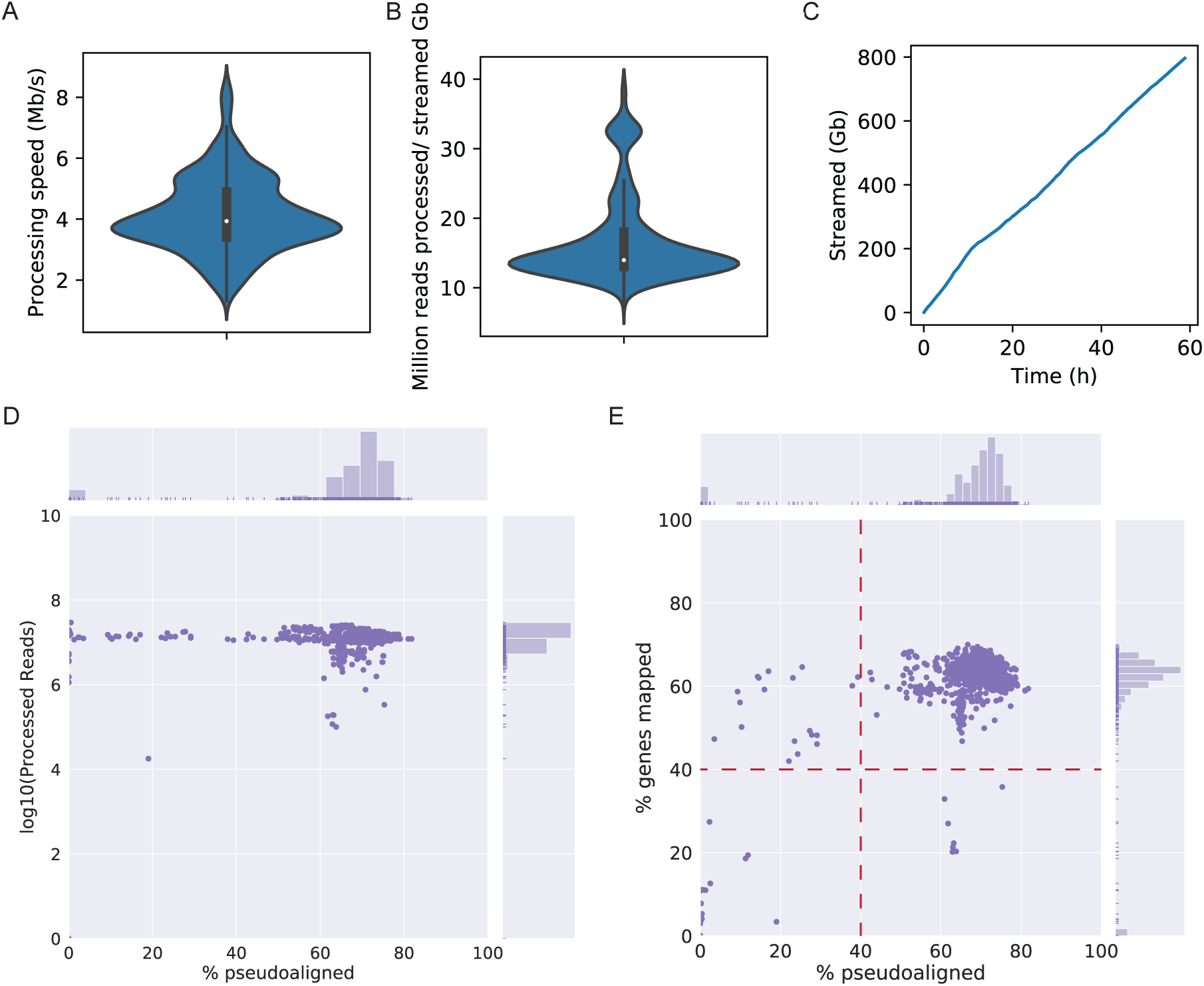
Summary of tobacco experiments streamed and processed by LSTrAP-Cloud. **(a)** The violin plot shows the speed of simultaneous streaming and processing of RNA-seq files by kallisto. **(b)** Amount of reads processed per gigabyte streamed. **(c)** Cumulative size (y-axis) of files streamed over time (x-axis). **(d)** The scatterplot shows the percentage of reads pseudoaligned to the CDS (x-axis) against the total number of streamed reads (y-axis). **(e)** The percentage of reads pseudoaligned to the CDS (x-axis) versus the percentage of genes with non-zero TPM values (y-axis). Red lines indicate cutoffs that were used to the select experiments for downstream analyses.

**Figure 3.**
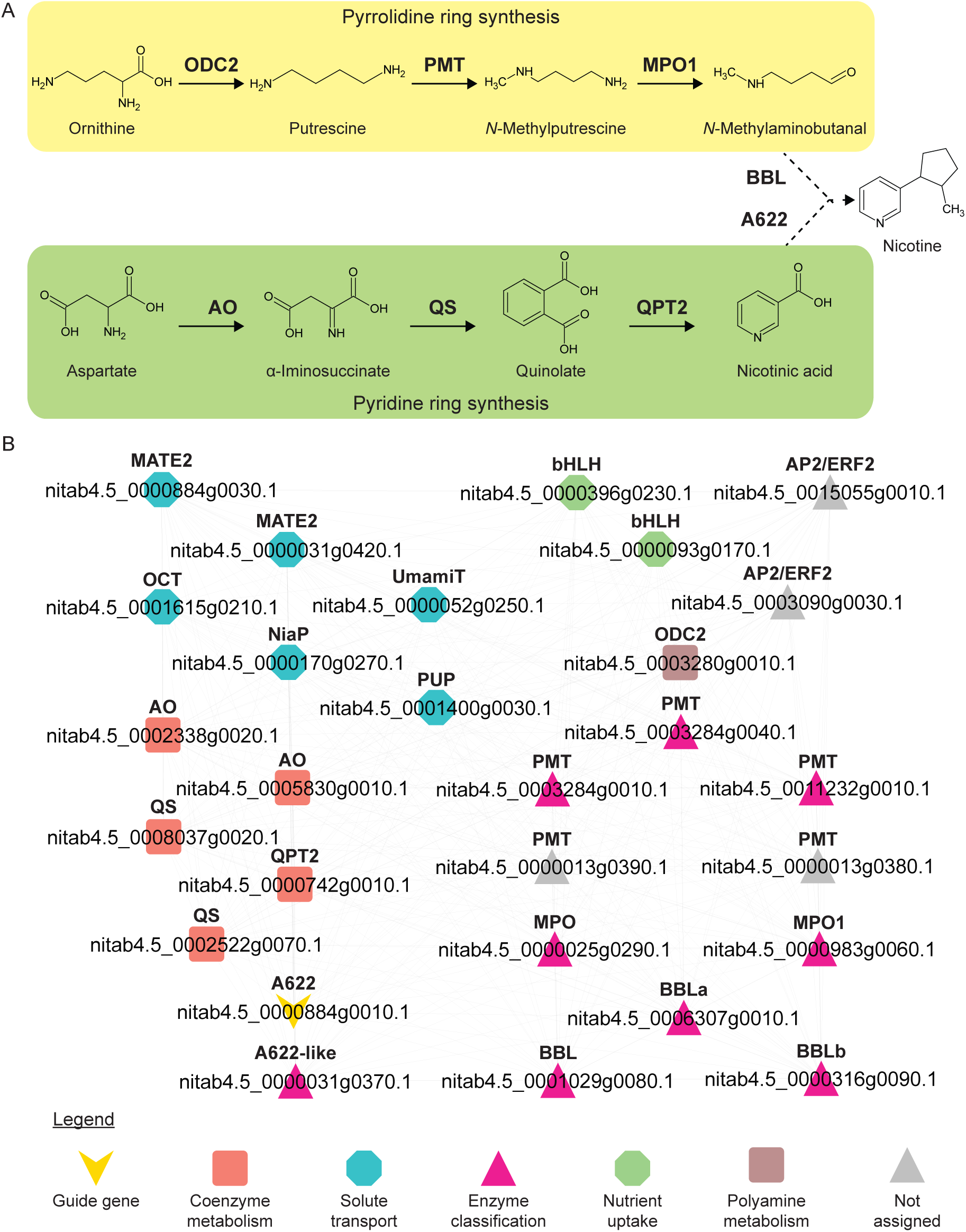
Nicotine biosynthesis. **(a)** Schematic of the nicotine biosynthesis pathway. Abbreviations for the enzymes are ODC2: ornithine decarboxylase 2, PMT: putrescine N-methyltransferase, MPO1: N-methylputrescine oxidase 1, BBL: berberine bridge enzyme-like proteins, AO: L-aspartate oxidase, QS: quinolinate synthase, QPT2: quinolinate phosphoribosyltransferase 2, A622: isoflavone-like oxidoreductase. **(b)** Co-expression network of A622 (Nitab4.5_0000884g0010.1). Abbreviations are AP2/ERF: APETALA 2/ethylene responsive factor, bHLH: basic helix-loop-helix, MATE2: multi antimicrobial extrusion family protein 2, OCT: organic cation transporter, NiaP: nicotinate transporter, PUP: purine uptake permease, and UmamiT: Usually Multiple Amino Acids Move In and Out Transporters. For brevity, only homologs of genes involved in nicotine biosynthesis as described in **(a)**, transporters and transcription factors are shown.

### 2.3. Identification of genes involved in nicotine biosynthesis

Nucleotide sequences of genes that reported to be involved in nicotine biosynthesis and transport (ODC1: AB031066; ODC2: AF233849; PMT: D28506; MPO1: AB289456; MPO2: AB289457; AO: XM_016633697, XR_001648132; QS: XM_016642986, XM_016638757; QPT1: AJ748262; QPT2: AJ748263; A622: D28505; BBLa: AB604219; BBLb: AM851017; BBLc: AB604220; BBLd: AB604221; MATE1: AB286963; MATE2: AB286962 and NUP1: GU174267) [33] were retrieved from NCBI. The gene IDs of these genes in the tobacco genome version Nitab-v4.5_cDNA_Edwards2017.fasta were identified through blast v2.6.0+ against the *N. tabacum* CDS. The function of the genes found in the network was further annotated using results from blast and Mercator (Table S3 and 5).

### 2.4. Relative expression of nicotine biosynthesis genes in 5 major organs

To obtain the relative expression of the genes shown in the co-expression network of *Nitab4.5_0000884g0010.1*, annotation of experiments were retrieved from NCBI SRA run selector (Table S1). Only wild-type and untreated experiments indicating leaf, flower, root, shoot and stem were selected for the analysis (Table S6). The median expression value of a gene in an organ was normalised with the highest median expression value of the gene across all organs.

## 3. Results

### 3.1. Implementation of gene co-expression pipeline on Google Colaboratory

The improvement in transcript estimation algorithms has greatly reduced the amount of time and resources required to estimate gene expression from RNA-sequencing data. Previously, we have demonstrated with the LSTrAP-Lite pipeline that analysis of large scale transcriptomic data was possible on a small computer costing less than 50 USD [24]. However, user friendliness of LSTrAP-Lite pipeline was still limited, as it runs in the linux terminal and ARM CPU architecture, which is not user-friendly to most biologists and not compatible with most software, respectively.

Here, we implemented a large scale transcriptomic analysis pipeline on Google Colaboratory, a free cloud computing platform that allows multiple users to easily share and deploy python code in a jupyter notebook environment. The pipeline (Figure 1), Large Scale Transcriptome Analysis Pipeline in Cloud (LSTrAP-Cloud), takes the CDS of the organism of interest, and streams the list of RNA-seq experiments specified by the user from ENA. After all files have been streamed, quality statistics from the download report are displayed on the notebook and summarised in plots. The plots allow the user to set an appropriate cutoff for experiments to be included in downstream analyses. Lastly, the pipeline generates and displays a co-expression network of the gene of interest, which can be downloaded as a PNG or JSON file. All outputs from the notebook are saved in a Google Drive account of the user, which is needed to run the notebook. We provide a user’s manual (Document S1) and SRA experiment list (Table S1), allowing the readers to replicate this analysis. All the files and data are conveniently stored on the user’s Google Drive.

The performance of the LSTrAP-Cloud was evaluated on the gene expression data of *Nicotiana tabacum*, a commercially important crop for the production of tobacco and an important model used in plant research. The first billion (109) bytes of all publicly available RNA sequencing experiments of *N. tabacum* were streamed from ENA and processed by kallisto. An average processing speed of 4 Mb/s was achieved (Figure 2a) and an average of 17 million reads were pseudoaligned per 1 Gb of data (Figure 2b). Overall, 796 Gb of data was streamed in 59 hours (Figure 2c), which is equivalent to 4 minutes per Gb. Quality statistics revealed that most experiments had more than 107 reads processed by kallisto, where 60-80% of the reads were pseudoaligned to the CDS (Figure 2d), and 50-70% of the CDS had a non-zero expression (Figure 2e). Under the assumption that most samples are of good quality, we chose RNA-seq experiments that had at least one million reads pseudoaligned to the *N. tabacum* CDS, 40% of streamed reads mapped to the CDS and at least 40% of genes with non-zero Transcripts Per Kilobase Million (TPM) expression (Figure 2e, Table S2). 962 experiments passed these conditions and were compiled into an TPM expression matrix, where genes were arranged in rows and experiments in columns.

### 3.2. Investigating the nicotine biosynthesis co-expression network

Nicotine is a toxic alkaloid produced by plants in the Solanaceae family to deter herbivores and a potent addictive substance. The synthesis of nicotine occurs in the roots and involves two precursors, a pyrrolidine (N-methyl-Δ^1^-pyrrolinium cation) and a pyridine (nicotinic acid) ring derived by a series of reactions from ornithine and aspartate, respectively ([34], Figure 3a). The rings are then combined to by the enzymes A622 (isoflavone reductase) and berberine bridge enzyme-like (BBL) to form nicotine. After synthesis in the roots, nicotine is sequestered out of the roots by multidrug and toxic compound extrusion (MATE) family transporters [35] and accumulated in organs that are highly prone to attack by herbivores, such as leaves [36].

Co-expression networks have been shown to be useful in the identification of enzymes, transcription factors and probable transporters of plant metabolites [11,20,24,37]. To demonstrate that this is also true in the case of nicotine biosynthesis, we retrieved the top 50 genes co-expressed with A622 (*Nitab4.5_0000884g0010.1*) (Figure 3b). The network revealed enzymes that are known to be involved in nicotine biosynthesis [aspartate oxidase (AO), quinolate synthase (QS), quinolinate phosphoribosyltransferase 2 (QPT2), ornithine decarboxylase 2 (ODC2), putrescine N-methytransferase (PMT), N-methylputrescine oxidase (MPO) and BBL] and transcription factors [APETALA 2/ ethylene responsive factor (AP2/ERF) and basic helix-loop-helix (bHLH)], which regulate nicotine biosynthesis [38]. In addition to the expected MATE2 transporters, other transporters such as organic cation transporter (OCT), nicotinate transporter (NiaP), Usually Multiple Amino Acids Move In and Out Transporter (UmamiT) and purine uptake permease (PUP) are also observed. To conclude, from the co-expression network of A622, we observed that more than half of the genes in the network (27 out of 50) are involved in nicotine biosynthesis. The remaining genes found in this co-expression network are excellent candidates for further functional analysis of their involvement in nicotine biosynthesis.

### 3.3. Expression analysis of genes related to nicotine biosynthesis in flower, leaf, root, shoot and stem

It is well established that the synthesis of nicotine occurs in the root, and numerous analyses have shown that secondary metabolism is under strong transcriptional control [7,24,39,40]. Hence, the expression of nicotine biosynthesis genes should be either specific or highly expressed in the roots. To confirm this, we calculated the median gene expression value of the genes found in the co-expression network (Figure 3) in roots, leaves, flowers, shoots and stems (Figure 4). As expected, all enzymes directly involved in nicotine biosynthesis are most expressed in roots and lowly expressed in other organs. Thus, we can conclude that the genes identified in the network of A622 are specifically expressed in the roots of *N. tabacum* are very likely to be involved in nicotine biosynthesis.

**Figure 4.**
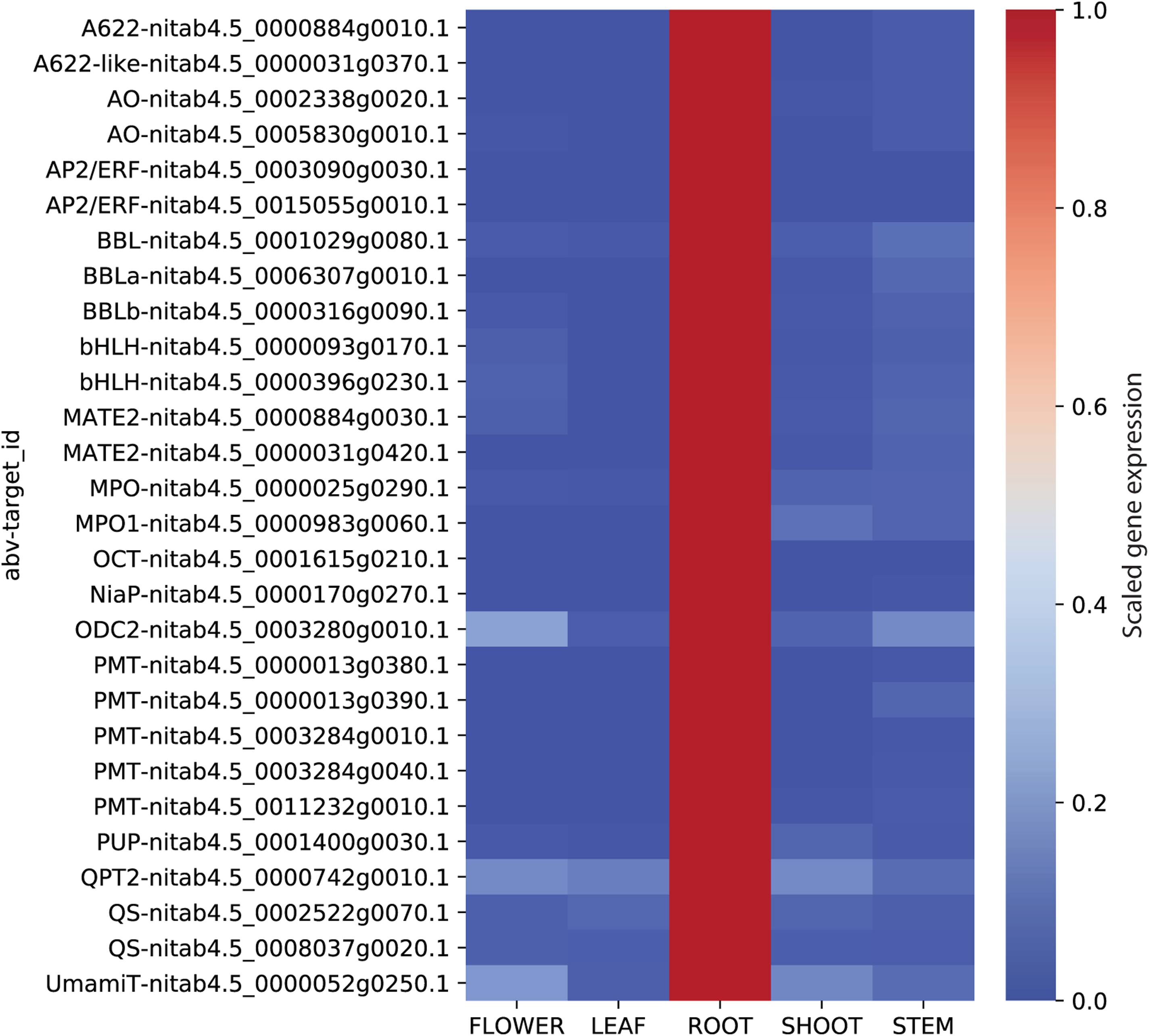
Relative expression of genes across major organs in *Nicotiana tabacum*. The expression values for each gene were scaled by dividing each row by the maximum value found in the row.

## 4. Discussion

As we sit on an expanding trove of data today, there is an immense amount of knowledge to be uncovered with the improvement in gene annotation and characterisation. Classical genomic approaches have allowed us to rapidly annotate genomes in silico based on sequence similarity to existing sequences. This approach has its limitations but can be greatly improved when the spatial and temporal expression of genes is taken into account.

In this study, we leveraged on the benefits of cloud computing and the user-friendliness of the Jupyter notebooks to implement a large-scale transcriptomic analysis pipeline, LSTrAP-Cloud (Figure 1). Using *N. tabacum* as an example (Figure 2), we showed that co-expression networks not only identified the enzymes involved in the metabolism of nicotine but also regulators and transporters that are found up- and down-stream of nicotine biosynthesis (Figures 3 and 4). The example of nicotine biosynthesis demonstrates that co-expression networks analysis is a valuable addition to sequence similarity-based approaches as it can infer modules of functionally related genes.

While the field of bioinformatics is advancing rapidly, it is important that biologists are also empowered with the tools and predictions available to bioinformaticians, as this can greatly shorten the amount of time required for gene characterisation through the identification of potential targets. The future of gene function prediction, however, will require a new generation of biologist equipped to tackle both wet and dry lab as sequencing data becomes available at a faster and larger scale [41].

## Supplementary Materials

The following are available online at http://www.mdpi.com/2073-4425/xx/1/5/s1, Figure S1: Complete co-expression network of A622 (*Nitab4.5_0000884g0010.1*), Table S1: SRA run table from NCBI SRA, Table S2: Download report of *N. tabacum* RNA-seq experiments streamed using LSTrAP-Cloud, Table S3: Annotation of *N. tabacum* CDS by Mercator, Table S4: Neighbourhood of A622, Table S5: BLAST output of genes involved in nicotine biosynthesis, Table S6: Summary of RNA-seq experiments included in the calculation of relative gene expression, Document S1: User manual for LSTrAP-Cloud

## Author Contributions

G.W. and T.Q. designed and established the pipeline. T.Q. implemented the pipeline and interpreted the results. T.Q. and M.M wrote the manuscript. M.M. conceptualised and supervised the project.

## Funding

This research was funded by Nanyang Technological University Start Up Grant, Singapore.

## Acknowledgments

We will like to thank Nanyang Technological University for funding and Lior Pachter for his useful discussions.

## Conflicts of Interest

The authors declare no conflict of interest.

## Abbreviations

The following abbreviations are used in this manuscript:

A622: isoflavone reductase
AO: aspartate oxidase
AP2/ERF: APETALA 2/ethylene responsive factor
BBL: berberine bridge enzyme-like
bHLH: basic helix-loop-helix
ENA: European Sequencing Archive
LSTrAP-Cloud: Large-Scale Transcriptomic Analysis Pipeline in Cloud
MATE2: multi antimicrobial extrusion family protein 2
MPO: N-methylputrescine oxidase
NiaP: nicotinate transporter
OCT: organic cation transporter
ODC2: ornithine decarboxylase 2
PMT: putrescine N-methytransferase
PUP: purine uptake permease
QPT2: quinolinate phosphoribosyltransferase
QS: quinolate synthase
RNA-seq: RNA sequencing
SRA: NCBI Sequence Read Archive
UmamiT: Usually Multiple Amino Acids Move In and Out Transporters

## Sample Availability

The expression matrix is found in the supplemental data and the pipeline is available at https://github.com/tqiaowen/LSTrAP-Cloud.

---

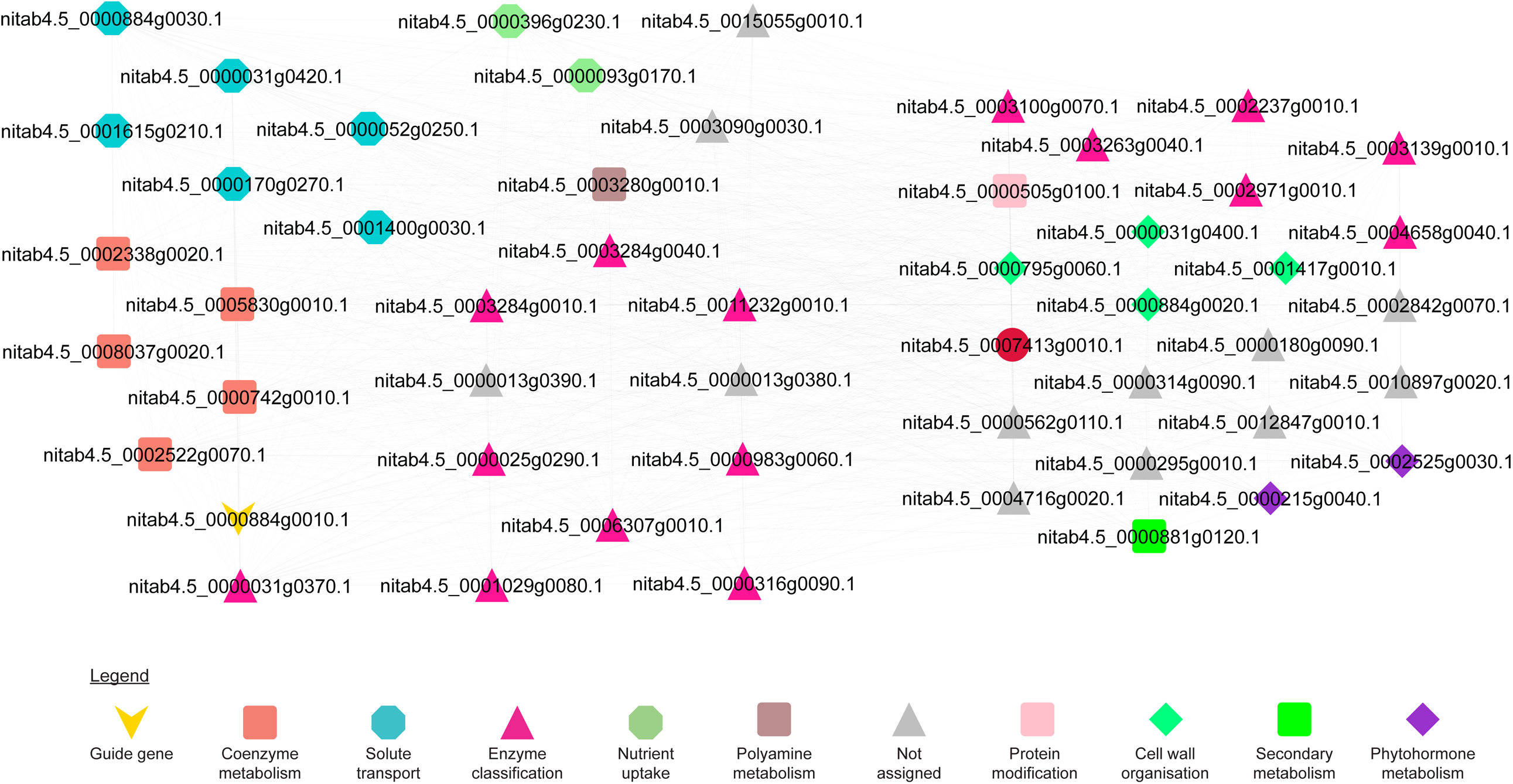

## References

1. Senol Cali, D.; Kim, J.S.; Ghose, S.; Alkan, C.; Mutlu, O. Nanopore sequencing technology and tools for genome assembly: computational analysis of the current state, bottlenecks and future directions. Briefings in Bioinformatics 2019, 20, 1542–1559. doi: 10.1093/bib/bby017.

2. Hansen, B.O.; Meyer, E.H.; Ferrari, C.; Vaid, N.; Movahedi, S.; Vandepoele, K.; Nikoloski, Z.; Mutwil, M. Ensemble gene function prediction database reveals genes important for complex I formation in Arabidopsis thaliana. New Phytologist 2018, 217, 1521–1534. doi: 10.1111/nph.14921.

3. Rhee, S.Y.; Mutwil, M. Towards revealing the functions of all genes in plants. Trends in Plant Science 2014, 19, 212–221. doi: 10.1016/j.tplants.2013.10.006.

4. Bouché, N.; Bouché, D. Arabidopsis gene knockout: phenotypes wanted. Current Opinion in Plant Biology 2001, 4, 111–117. doi: 10.1016/S1369-5266(00)00145-X.

5. Zhang, X.; Henriques, R.; Lin, S.S.; Niu, Q.W.; Chua, N.H. Agrobacterium-mediated transformation of Arabidopsis thaliana using the floral dip method. Nature Protocols 2006, 1, 641–646. doi: 10.1038/nprot.2006.97.

6. Ruprecht, C.; Vaid, N.; Proost, S.; Persson, S.; Mutwil, M. Beyond Genomics: Studying Evolution with Gene Coexpression Networks. Trends in Plant Science 2017, 22. doi: 10.1016/j.tplants.2016.12.011.

7. Ruprecht, C.; Mendrinna, A.; Tohge, T.; Sampathkumar, A.; Klie, S.; Fernie, A.R.; Nikoloski, Z.; Persson, S.; Mutwil, M. FamNet: A framework to identify multiplied modules driving pathway diversification in plants. Plant Physiology 2016, 170, 1878–1894. doi: 10.1104/pp.15.01281.

8. Usadel, B.; Obayashi, T.; Mutwil, M.; Giorgi, F.M.; Bassel, G.W.; Tanimoto, M.; Chow, A.; Steinhauser, D.; Persson, S.; Provart, N.J. Co-expression tools for plant biology: Opportunities for hypothesis generation and caveats. Plant, Cell and Environment 2009, 32, 1633–1651. doi: 10.1111/j.1365-3040.2009.02040.x.

9. Lee, I.; Ambaru, B.; Thakkar, P.; Marcotte, E.M.; Rhee, S.Y. Rational association of genes with traits using a genome-scale gene network for Arabidopsis thaliana. Nature Biotechnology 2010, 28, 149–156. doi: 10.1038/nbt.1603.

10. Hansen, B.O.; Vaid, N.; Musialak-Lange, M.; Janowski, M.; Mutwil, M. Elucidating gene function and function evolution through comparison of co-expression networks of plants. Frontiers in Plant Science 2014, 5, 1–9. doi: 10.3389/fpls.2014.00394.

11. Proost, S.; Mutwil, M. Tools of the trade: Studying molecular networks in plants. Current Opinion in Plant Biology 2016, 30, 130–140. doi: 10.1016/j.pbi.2016.02.010.

12. Takabayashi, A.; Ishikawa, N.; Obayashi, T.; Ishida, S.; Obokata, J.; Endo, T.; Sato, F. Three novel subunits of Arabidopsis chloroplastic NAD(P)H dehydrogenase identified by bioinformatic and reverse genetic approaches. Plant Journal 2009, 57, 207–219. doi: 10.1111/j.1365-313X.2008.03680.x.

13. Takahashi, N.; Lammens, T.; Boudolf, V.; Maes, S.; Yoshizumi, T.; De Jaeger, G.; Witters, E.; Inzé, D.; De Veylder, L. The DNA replication checkpoint aids survival of plants deficient in the novel replisome factor ETG1. The EMBO journal 2008, 27, 1840–1851. doi: 10.1038/emboj.2008.107.

14. Stuart, J.M.; Segal, E.; Koller, D.; Kim, S.K. A Gene-Coexpression Network for Global Discovery of Conserved Genetic Modules. Science 2003, 302, 249–255. doi: 10.1126/science.1087447.

15. Yu, H.; Luscombe, N.M.; Qian, J.; Gerstein, M. Genomic analysis of gene expression relationships in transcriptional regulatory networks, 2003. doi: 10.1016/S0168-9525(03)00175-6.

16. Jiménez-Gómez, J.M.; Wallace, A.D.; Maloof, J.N. Network analysis identifies ELF3 as a QTL for the shade avoidance response in arabidopsis. PLoS Genetics 2010, 6. doi: 10.1371/journal.pgen.1001100.

17. Persson, S.; Wei, H.; Milne, J.; Page, G.P.; Somerville, C.R. Identification of genes required for cellulose synthesis by regression analysis of public microarray data sets. Proceedings of the National Academy of Sciences of the United States of America 2005, 102, 8633–8638. doi: 10.1073/pnas.0503392102.

18. Itkin, M.; Heinig, U.; Tzfadia, O.; Bhide, a.J.; Shinde, B.; Cardenas, P.D.; Bocobza, S.E.; Unger, T.; Malitsky, S.; Finkers, R.; Tikunov, Y.; Bovy, A.; Chikate, Y.; Singh, P.; Rogachev, I.; Beekwilder, J.; Giri, A.P.; Aharoni, A. Biosynthesis of antinutritional alkaloids in solanaceous crops is mediated by clustered genes. Science 2013, 341, 175–179. doi: 10.1126/science.1240230.

19. Proost, S.; Mutwil, M. PlaNet: Comparative Co-Expression Network Analyses for Plants. In Methods in Molecular Biology; van Dijk, A.D.J., Ed.; Springer New York: New York, NY, 2017; Vol. 1533, pp. 213–227. doi: 10.1007/978-1-4939-6658-5_12.

20. Sibout, R.; Proost, S.; Hansen, B.O.; Vaid, N.; Giorgi, F.M.; Ho-Yue-Kuang, S.; Legée, F.; Cézart, L.; Bouchabké-Coussa, O.; Soulhat, C.; Provart, N.; Pasha, A.; Le Bris, P.; Roujol, D.; Hofte, H.; Jamet, E.; Lapierre, C.; Persson, S.; Mutwil, M. Expression atlas and comparative coexpression network analyses reveal important genes involved in the formation of lignified cell wall in Brachypodium distachyon. New Phytologist 2017, 215, 1009–1025. doi: 10.1111/nph.14635.

21. Alejandro, S.; Lee, Y.Y.; Tohge, T.; Sudre, D.; Osorio, S.; Park, J.; Bovet, L.; Lee, Y.Y.; Geldner, N.; Fernie, A.R.; Martinoia, E. AtABCG29 is a monolignol transporter involved in lignin biosynthesis. Current Biology 2012, 22, 1207–1212. doi: 10.1016/j.cub.2012.04.064.

22. Bray, N.L.; Pimentel, H.; Melsted, P.; Pachter, L. Near-optimal probabilistic RNA-seq quantification. Nature Biotechnology 2016, 34, 525–527, [1505.02710]. doi: 10.1038/nbt.3519.

23. Patro, R.; Duggal, G.; Love, M.I.; Irizarry, R.A.; Kingsford, C. Salmon provides fast and bias-aware quantification of transcript expression. Nature methods 2017, 14, 417–419. doi: 10.1038/nmeth.4197.

24. Tan, Q.W.; Mutwil, M. Inferring biosynthetic and gene regulatory networks from Artemisia annua RNA sequencing data on a credit card-sized ARM computer. Biochimica et Biophysica Acta (BBA) - Gene Regulatory Mechanisms 2019, p. 194429. doi: 10.1016/j.bbagrm.2019.194429.

25. Kohen, R.; Barlev, J.; Hornung, G.; Stelzer, G.; Feldmesser, E.; Kogan, K.; Safran, M.; Leshkowitz, D. UTAP: User-friendly Transcriptome Analysis Pipeline. BMC Bioinformatics 2019, 20, 154. doi: 10.1186/s12859-019-2728-2.

26. Vaneechoutte, D.; Vandepoele, K. Curse: building expression atlases and co-expression networks from public RNA-Seq data. Bioinformatics 2019, 35, 2880–2881. doi: 10.1093/bioinformatics/bty1052.

27. Proost, S.; Krawczyk, A.; Mutwil, M. LSTrAP: Efficiently combining RNA sequencing data into co-expression networks. BMC Bioinformatics 2017, 18. doi: 10.1186/s12859-017-1861-z.

28. Leinonen, R.; Akhtar, R.; Birney, E.; Bower, L.; Cerdeno-Tárraga, A.; Cheng, Y.; Cleland, I.; Faruque, N.; Goodgame, N.; Gibson, R.; Hoad, G.; Jang, M.; Pakseresht, N.; Plaister, S.; Radhakrishnan, R.; Reddy, K.; Sobhany, S.; Ten Hoopen, P.; Vaughan, R.; Zalunin, V.; Cochrane, G. The European Nucleotide Archive. Nucleic acids research 2011, 39, D28–D31. doi: 10.1093/nar/gkq967.

29. Edwards, K.D.; Fernandez-Pozo, N.; Drake-Stowe, K.; Humphry, M.; Evans, A.D.; Bombarely, A.; Allen, F.; Hurst, R.; White, B.; Kernodle, S.P.; Bromley, J.R.; Sanchez-Tamburrino, J.P.; Lewis, R.S.; Mueller, L.A. A reference genome for Nicotiana tabacum enables map-based cloning of homeologous loci implicated in nitrogen utilization efficiency. BMC Genomics 2017, 18, 448. doi: 10.1186/s12864-017-3791-6.

30. Fernandez-Pozo, N.; Menda, N.; Edwards, J.D.; Saha, S.; Tecle, I.Y.; Strickler, S.R.; Bombarely, A.; Fisher-York, T.; Pujar, A.; Foerster, H.; Yan, A.; Mueller, L.A. The Sol Genomics Network (SGN)—from genotype to phenotype to breeding. Nucleic Acids Research 2015, 43, D1036–D1041. doi: 10.1093/nar/gku1195.

31. Franz, M.; Lopes, C.T.; Huck, G.; Dong, Y.; Sumer, O.; Bader, G.D. Cytoscape.js: A graph theory library for visualisation and analysis. Bioinformatics 2015, 32, 309–311. doi: 10.1093/bioinformatics/btv557.

32. Schwacke, R.; Ponce-Soto, G.Y.; Krause, K.; Bolger, A.M.; Arsova, B.; Hallab, A.; Gruden, K.; Stitt, M.; Bolger, M.E.; Usadel, B. MapMan4: A Refined Protein Classification and Annotation Framework Applicable to Multi-Omics Data Analysis. Molecular Plant 2019, 12, 879–892. doi: 10.1016/j.molp.2019.01.003.

33. Kajikawa, M.; Sierro, N.; Kawaguchi, H.; Bakaher, N.; Ivanov, N.V.; Hashimoto, T.; Shoji, T. Genomic Insights into the Evolution of the Nicotine Biosynthesis Pathway in Tobacco. Plant Physiology 2017, 174, 999–1011. doi: 10.1104/pp.17.00070.

34. Xu, S.; Brockmöller, T.; Navarro-Quezada, A.; Kuhl, H.; Gase, K.; Ling, Z.; Zhou, W.; Kreitzer, C.; Stanke, M.; Tang, H.; Lyons, E.; Pandey, P.; Pandey, S.P.; Timmermann, B.; Gaquerel, E.; Baldwin, I.T. Wild tobacco genomes reveal the evolution of nicotine biosynthesis. Proceedings of the National Academy of Sciences 2017, 114, 6133–6138. doi: 10.1073/pnas.1700073114.

35. Shoji, T.; Inai, K.; Yazaki, Y.; Sato, Y.; Takase, H.; Shitan, N.; Yazaki, K.; Goto, Y.; Toyooka, K.; Matsuoka, K.; Hashimoto, T. Multidrug and Toxic Compound Extrusion-Type Transporters Implicated in Vacuolar Sequestration of Nicotine in Tobacco Roots. Plant Physiology 2009, 149, 708–718. doi: 10.1104/pp.108.132811.

36. Baldwin, I.T. An Ecologically Motivated Analysis of Plant-Herbivore Interactions in Native Tobacco. Plant Physiology 2001, 127, 1449–1458. doi: 10.1104/pp.010762.

37. Ruprecht, C.; Mutwil, M.; Saxe, F.; Eder, M.; Nikoloski, Z.; Persson, S. Large-Scale Co-Expression Approach to Dissect Secondary Cell Wall Formation Across Plant Species. Frontiers in Plant Science 2011, 2, 1–13. doi: 10.3389/fpls.2011.00023.

38. Liu, H.; Kotova, T.I.; Timko, M.P. Increased Leaf Nicotine Content by Targeting Transcription Factor Gene Expression in Commercial Flue-Cured Tobacco (Nicotiana tabacum L.). Genes 2019, 10, 930. doi: 10.3390/genes10110930.

39. Mutwil, M.; Klie, S.; Tohge, T.; Giorgi, F.M.; Wilkins, O.; Campbell, M.M.; Fernie, A.R.; Usadel, B.; Nikoloski, Z.; Persson, S. PlaNet: Combined Sequence and Expression Comparisons across Plant Networks Derived from Seven Species. The Plant Cell 2011, 23, 895–910. doi: 10.1105/tpc.111.083667.

40. Ferrari, C.; Shivhare, D.; Hansen, B.O.; Pasha, A.; Esteban, E.; Provart, N.J.; Kragler, F.; Fernie, A.R.; Tohge, T.; Mutwil, M. Expression Atlas of Selaginella moellendorffii Provides Insights into the Evolution of Vasculature, Secondary Metabolism, and Roots. The Plant Cell 2020, p. tpc.00780.2019. doi: 10.1105/tpc.19.00780.

41. Friesner, J.; Assmann, S.M.; Bastow, R.; Bailey-Serres, J.; Beynon, J.; Brendel, V.; Buell, C.R.; Bucksch, A.; Busch, W.; Demura, T.; Dinneny, J.R.; Doherty, C.J.; Eveland, A.L.; Falter-Braun, P.; Gehan, M.A.; Gonzales, M.; Grotewold, E.; Gutierrez, R.; Kramer, U.; Krouk, G.; Ma, S.; Markelz, R.J.C.; Megraw, M.; Meyers, B.C.; Murray, J.A.H.; Provart, N.J.; Rhee, S.; Smith, R.; Spalding, E.P.; Taylor, C.; Teal, T.K.; Torii, K.U.; Town, C.; Vaughn, M.; Vierstra, R.; Ware, D.; Wilkins, O.; Williams, C.; Brady, S.M. The Next Generation of Training for Arabidopsis Researchers: Bioinformatics and Quantitative Biology. Plant Physiology 2017, 175, 1499–1509. doi: 10.1104/pp.17.01490.

