## Supplementary material for "LSTrAP-Cloud: A User-friendly Cloud Computing Pipeline to Infer Co-functional and Regulatory Networks": Document S1

### LSTrAP-Cloud

Large-Scale Transcriptome Analysis Pipeline on Cloud
This repository is built upon wirriamm/CoNeGC

#### What is LSTrAP-Cloud?

LSTrAP-Cloud is a pipeline designed for building co-expression networks from RNA-seq data (fastq files from ENA) on Goolge Colaboratory (Colab). Leveraging on the user-friendliness of the Colab interface, LSTrAP-Cloud allows users to analyse large scale transcriptome data without having to access the linux terminal, making it accessible to both bioinformaticians and biologist. To get started, we have provided a tutorial based on example data found here. While the pipeline was designed for plants, we have also made the script compatible with non-plant organisms.

#### Changelog

#### Tutorial sections


1. Preparation of Google Drive Account
2. Setting up Google Colaboratory
   2.1 Opening the pipeline on Google Colab
   2.2 Running code in Cells
   2.3 Connecting to your Google Drive account
3. Streaming RNA-seq data
4. Generating Neighbourhood and Network Files
   4.1 User input of variables
   4.2 Setting the threshold for acceptable RunIDs
   4.3 Creating the gene co-expression network


#### Acknowledgements


LSTrAP-Cloud will not have been possible without the various open-source projects.


#### Contact

Issues and feedback can be submitted through GitHub or to Qiao Wen Tan.

### Tutorial

Example files are provided to help you get started with the pipeline.

#### 1. Preparation of Google Drive Account

Please ensure sufficient storage space of more than 1 GB. Space requirement varies with the organism and number of experiments you wish to analyse.

With your Google Drive account, create a directory containing the following files:

| File | Remarks |
|---|---|
| runid.txt | Contains list of RunIDs to be streamed from ENA. Example here |
| CDS.fastq | CDS file of the organism RNA-seq experiments are to be mapped to. gz compressed files are also accepted. The CDS of *N. tabacum* (Nitab-v4.5_cDNA_Edwards2017.fasta) can be downloaded at SolGenomics |

#### 2. Setting up Google Colaboratory

##### 2.1 Opening the pipeline on Google Colab

There are two ways to open the notebook.
**Method 1: Opening in Colab** (File > Open Notebook > GitHub)

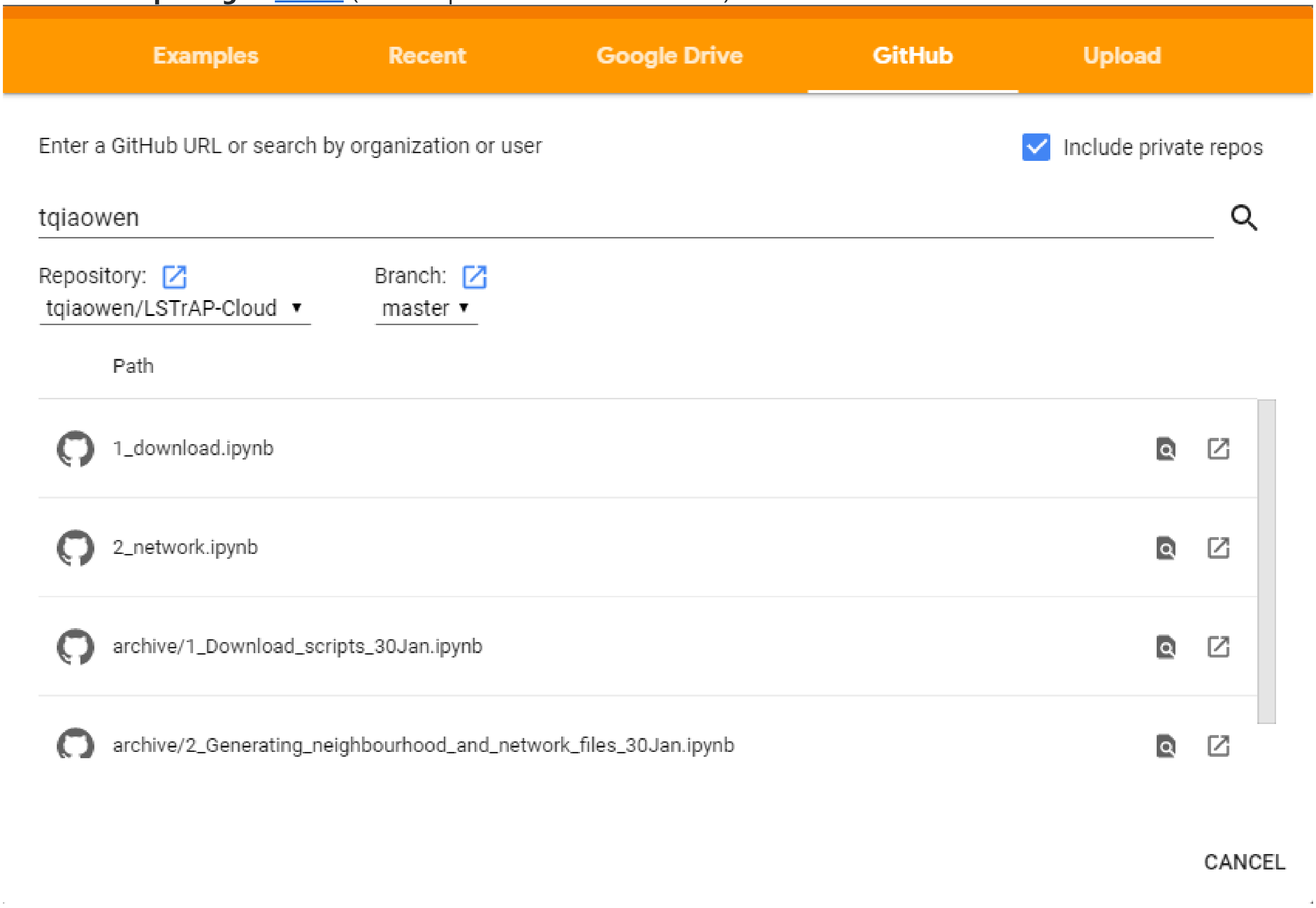

**Method 2: Opening through the [LSTrAP-Cloud repository]** (Click on 'Open in Colab')

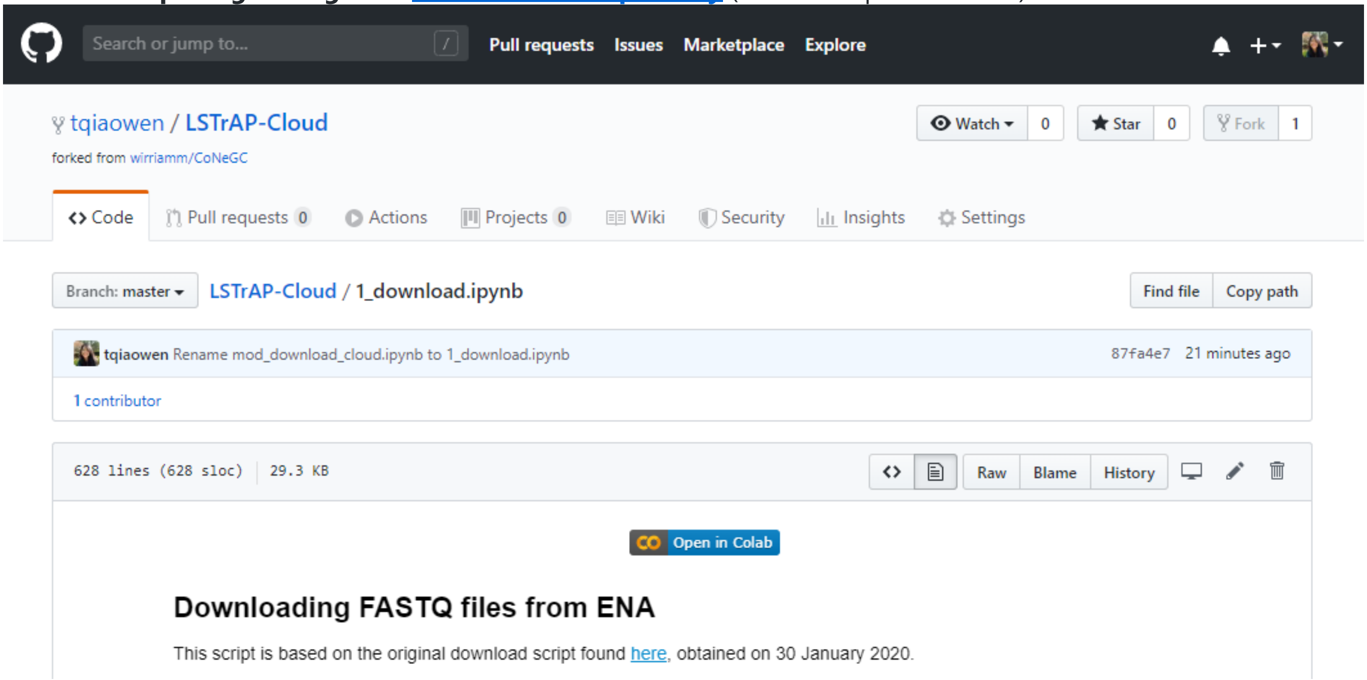


#### 2.2 Running code in cells

A cell can be run by clicking on the play button at the top left of each cell. The following options can be used to run multiple cells by clicking from the menu bar, or by hotkeys as specified in parentheses:

- Runtime > Run all (Crtl+F9): Runs all cells in the notebook
- Runtime > Run before (Crtl+F8): Runs all cells before the cell in focus
- Runtime > Run after (Crtl+F10): Runs all cells after the cell in focus Tip: To prevent the notebook from going idle while the script is running, a javascript code can be implemented in the web browser. Open the browser's javascript console and paste the following code and hit enter:

```
function ClickConnect(){
console.log("Working");
document.querySelector("colab-toolbar-button#connect").click()
}
setInterval(ClickConnect,60000)
```

#### 2.3 Connecting to your Google Drive account

To connect your Google Drive account to Colab, run the first cell (Cells 1.1 and 2.1 for 1_download.ipynb and 2_network.ipynb) and enter the authorisation code.

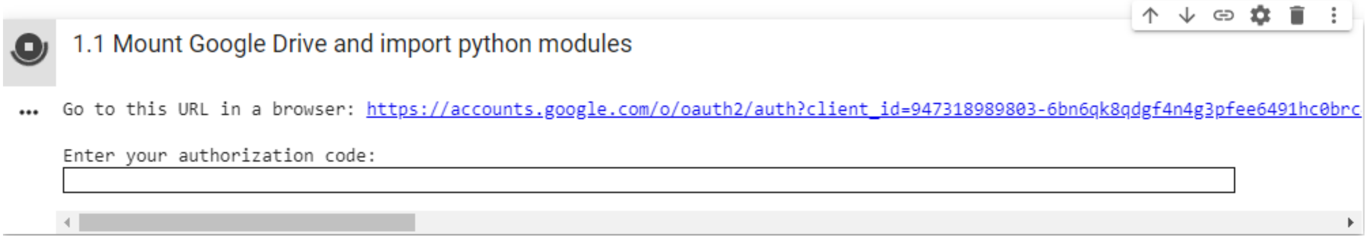


After mounting your Google Drive, do save a copy of the notebook to your drive (File > Save a copy in Drive)!

#### 3. Streaming RNA-seq data

Before running the rest of the cells, the notebook requires some information to be filled up under cells 1.2. After filling up the cell, the rest of the cells can be executed. **Note!**

- Include file extensions (eg. '.txt', '.tsv') in the file names
- Ensure that files are already saved in the Google Drive folder specified
- Avoid any whitespace in file names
- For new download, select 'A. Start fresh run'. The date initiated section can be ignored
- To continue from previous download due to disconnected runtime (can happen when attempting to download large amounts of experiments), select 'B. Continue with previous run' and select the date when the download was
initiated.

▾ User input of variables

[ ] 1.2 Input form

1) Enter the absolute path for the folder you are going to work with. Tip: After mounting your Google Drive, right click on the file and copy path.

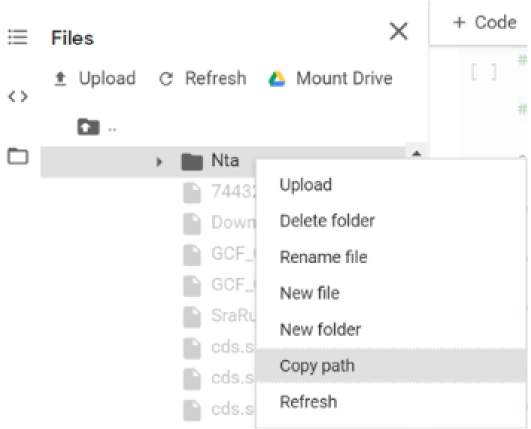


**in_dir:** " Insert text here "

2) Enter the species name. Eg. *Nicotiana tabacum* can be shortened to **Nta**

**species_name:** " Insert text here "

Expected outputs

| File | Remarks |
|---|---|
| index_file | Index file created by `kallisto index` based on the CDS provided |
| Kallisto output folders | Folders containing outputs generated by `kallisto quant` |
| Download_report.txt | Tab separated file summarising the status of download, amount of data downloaded, amount of time taken for kallisto streaming and a statistics from kallisto for each RunID. The file can be opened in Microsoft Excel. |

#### 4. Generating Neighbourhood and Network Files

This part of the tutorial will require you to use the second notebook. Refer to section 2.1 Opening the pipeline on Google Colab on how to do it. Do save a copy of the notebook to your Google Drive account after mounting your google drive.

Expected outputs

| File | Remarks |
|---|---|
| tpm.txt | Gene expression matrix generated based on the threshold provided |
| neighbourhood_file.txt | Contains information about the neighbourhood of the gene of interest such as geneID, PCC value and descriptions from mercator |
| network_file.txt | Contains information between gene pairs and their corresponding PCC values. Compatible with cytoscape desktop |
| gene_network.html | This file can be open standalone in a brower (tested on Chrome) and is the same file used to display the network in this notebook. |

##### 4.1 User input of variables

Similar to section 3. Streaming RNA-seq data, the fields in cell 2.2 should be filled with the respective directory or file paths which can be easily obtained by a right click on the directory/file and selecting 'Copy path'.

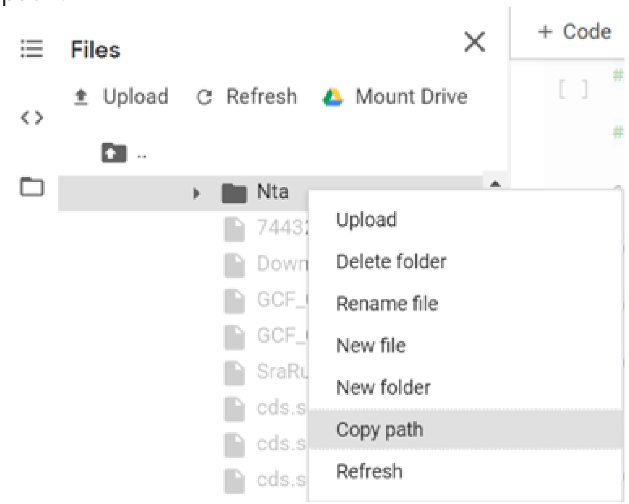


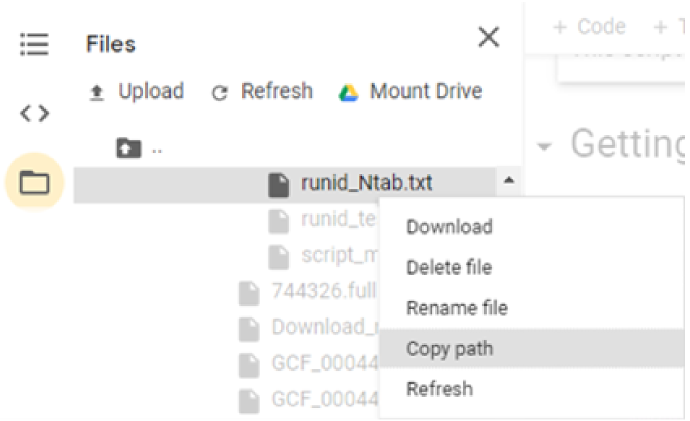


After filling up cell 2.2, run cells 2.2 to 2.6 to display the quality control table and scatter plots.

Note!

- If you are using a non-plant organism, please provide a file based on the mercator format. Gene identifiers should be in **lowercase**.
- An example of the mercator output for *N. tabacum* is also provided.

##### 4.2 Setting the threshold for acceptable RunIDs

After reviewing the quality control table and scatter plots, adjust the sliders in '2.7 Determine Quality Control Cutoff' to select the desired threshold levels. After adjusting the sliders, run cells 2.7 and 2.8 to extract the selected experiments and compile the gene expression matrix.

2.7 Determine Quality Control Cutoff

Please assess the quality of the downloaded files using the tables and graphs above. By adjusting the sliders below, only RunIDs (experiments) with values above the threshold will be selected for further analysis.

**p_pseudoaligned_threshold:** 40

Enter the $\log_{10}$ of reads processed instead of absolute number.

**reads_pseudoalinged_threshold:** 6

**p_genes_mapped_threshold:** 50

##### 4.3 Creating the gene co-expression network

After the gene expression matrix has been created, adjust the parameters in cell '2.9 Network options'. After adjusting the variables, run all the cells from 2.9 to 2.14.

| Variable | Remarks |
|---|---|
| goi | The gene identifier provided should be identical to that of the CDS file |
| cutoff | Pearson Correlation Coefficient cutoff to be used |
| neighbourhood_size | Number of neighbours that the network should contain excluding the gene of interest |
| is_a_plant | To indicate if the organism used for analysis is a plant |

[ ] 2.9 Network options

Description of gene to be analysed (must be the same as the one found in the CDS file)
*Eg. "lcl|NW_015787229.1_cds_XP_016497259.1_1"*

**goi:** " Insert text here "

**cutoff:** 0.7

**neighbourhood_size:** 51

Please uncheck if your organism is not a plant.

**is_a_plant:** ☑

The PNG and JSON format of the network can be downloaded by clicking on the links. The JSON file can be opened in Cytoscape desktop for further modifications to the network. The legend of the network and information regarding the genes in the network can be found in cells '2.13 Legend for nodes based on Mapman Bins:' and '2.14 Details of Genes in Network' respectively.

[ ] 2.12 Display Network

Click to download as:
Image | JSON

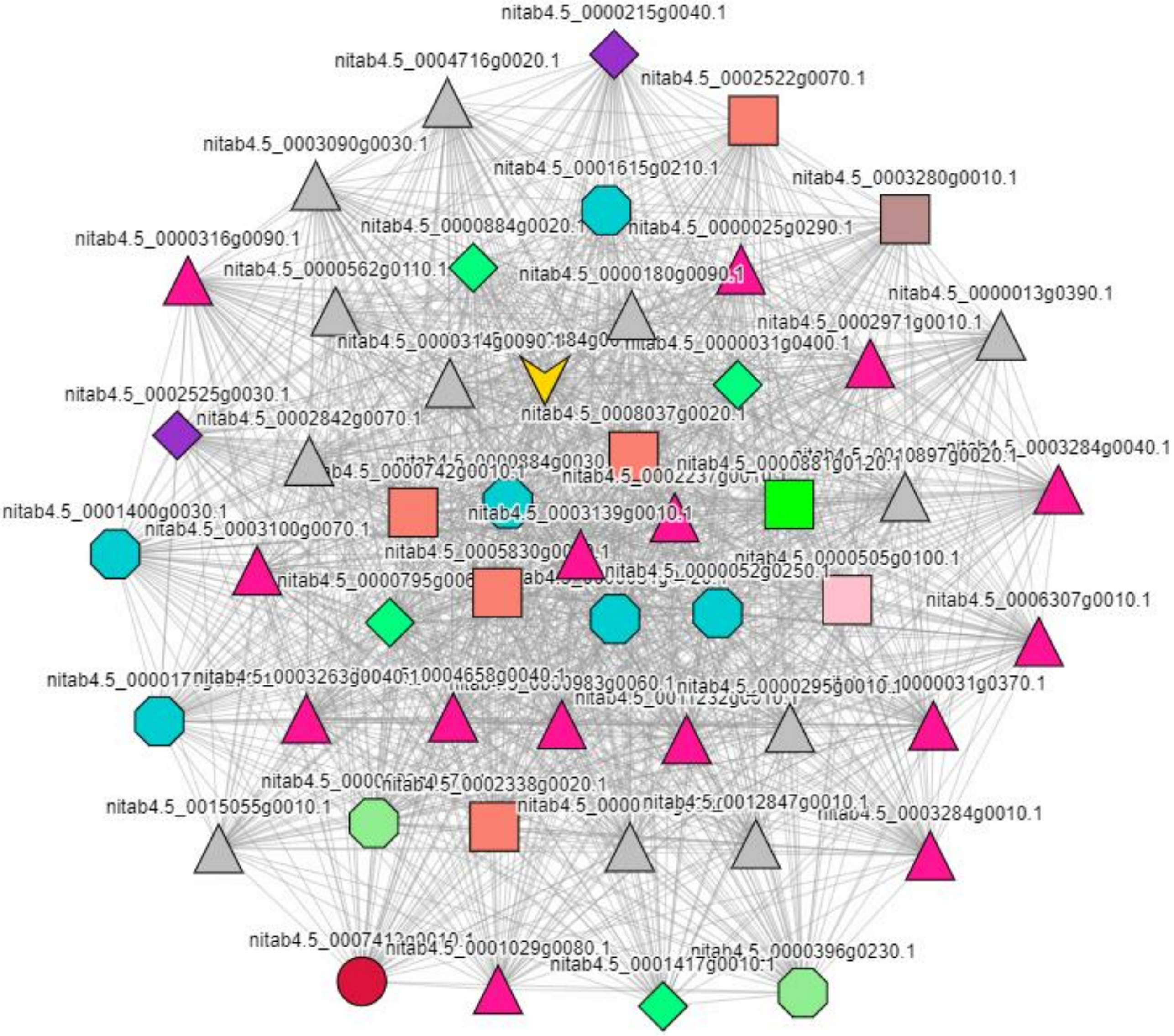
